# Action intentions result in the task-specific integration of object features

**DOI:** 10.1101/2025.04.28.651066

**Authors:** Nina Lee, Matthias Niemeier

## Abstract

Theories of object-based attention suggest that attending to an object binds its features together. Yet, there is a growing body of work to suggest that the intention to grasp an object can alter the representation of features such that they are separately represented during different stages of motor planning and execution, whereas some object features such as shape and size might form integrated representations when afforded by motor control. However, it remains untested whether these features were integrated as an outcome of the requirements of grasping motor control, or due to attention towards the object in general. Therefore, here we investigated how task-relevancy modulates the integration of grasp-relevant object features. To this end, we recorded electroencephalography while human participants grasped or reached for objects that varied in their orientation and size. Using multivariate analyses, we found a superadditive integration of object orientation and size during action planning for grasping but not reaching. These integrated representations likely facilitated the calculation of stable grasp points as further evidenced by the representations of grasp-specific visual size and grip size emerging at similar times. Our results provide novel insights into the vital role of action intention on cognitive representations in the human brain.

## Introduction

Attention directed to objects can result in its features being made simultaneously available for report (Duncan, 1984), or its features being bound together in an object file (Kahnemann et al., 1992). Together, theories of object-based attention suggest that the act of paying attention to an object should result in the conjunction of its features.

However, recent work in our lab has revealed that intending to grasp an object, compared to intending to simply reach for it, leads to differences in how grasp-relevant task features are represented in the brain (Lee et al., 2024). Specifically, we have found that representations for object shape, and for visual cues of weight form at different grasping phases, namely action planning and execution (Lee et al., 2024; see also Klein et al., 2023). This is consistent with the ability of sensorimotor control to compute task-relevant features in isolation (Ganel & Goodale, 2003; Nashed et al., 2012).

Nevertheless, in a grasping study, Guo and Niemeier (2024) found evidence that participants formed integrated representations of the shape and the size of the object after it became visible. One possible interpretation is that this integration reflects the calculation of grasp points on an object which requires considering both shape and size together (Blake, 1995) consistent with the idea that feature integration occurs when afforded by motor control. Alternatively however, integrated representations of the shape and size of the object might arise regardless of task as soon as attention is directed towards the object (Duncan, 1984; Kahnemann et al., 1992).

Thus, in the present EEG experiment, we investigated whether the intention to either grasp or reach for an object determines whether grasp-relevant object features are being integrated – or not. Further, we aimed to clarify the nature of size representations involved in the integrated representations of shape and size as Guo and Niemeier (2024) confounded object size with grip size (i.e., they used small and large objects that afforded small vs. large grasps).

To this end, human participants either grasped and pulled on an object or reached for an object to touch it with their knuckle (subsequently referred to as “knuckling”). Furthermore, we used objects that disentangled object size and grip size. That is, small and large objects all had the same cross shape with one long and one short axis but were presented in either a horizontal or vertical orientation. Participants grasped all objects horizontally and were thus required to perform the same sized grip on large vertical and small horizontal objects. We hypothesized that object orientation and grip size would form integrated representations, and that the integration would occur while participants planned to make grasp movements, not while they planned knuckling movements. Further, we expected to find representations of grip size and visual object size that occurred at similar times.

## Methods

### Participants

Fifteen undergraduate and graduate students (12 female; mean age: 21.3) from the University of Toronto participated in the experiment and were compensated $15/h. All students gave their written and informed consent prior to participating and all procedures were approved by the Human Participants Review Sub-Committee of the University of Toronto. All participants were screened to have normal or corrected-to-normal vision and right-hand dominance (Oldfield, 1971).

### Stimuli and Apparatus

Participants sat in a dark room at a table with a button box in front of a wooden mount that carried a shutter glass. The shutter glass permitted the viewing of target stimuli that were white 3D-printed objects with the same thickness (20.0 mm) and the same rounded cross shape (Fig. 1A). A black dot (30.0 mm across) was placed in the middle of each object to minimize the spread of laser light and to serve as a reach target. Objects were of two sizes: the large objects measured 67.5 mm by 45.0 mm along their secondary axis and were created by appending half-circles, 22.5 mm across, on each side of a 45.0 by 22.5 mm rectangle; the small objects measured 45.0 mm by 30.0 mm and were created by appending circles 15.0 mm in diameter to a 30.0 by 15.0 mm rectangle. Thus, large objects had a front face that was 55.6% larger in area than that of the small objects. These objects were then mounted such that either the longer or shorter axis was upright, giving two possible object orientations. Objects were mounted on rods in the back that were fitted into shafts boring into the centre of a frame measuring 170.0 by 105.0 mm, and four of these frames created the four sides of a motorized turntable. The apparatus was designed so that participants could easily grasp and pull the object on the front side and then quickly push the object back in place. The entire turntable was spray painted black to reduce the visual presence of the background and increase contrast between the background and the white objects (Fig. 1B).

**Figure 1.**
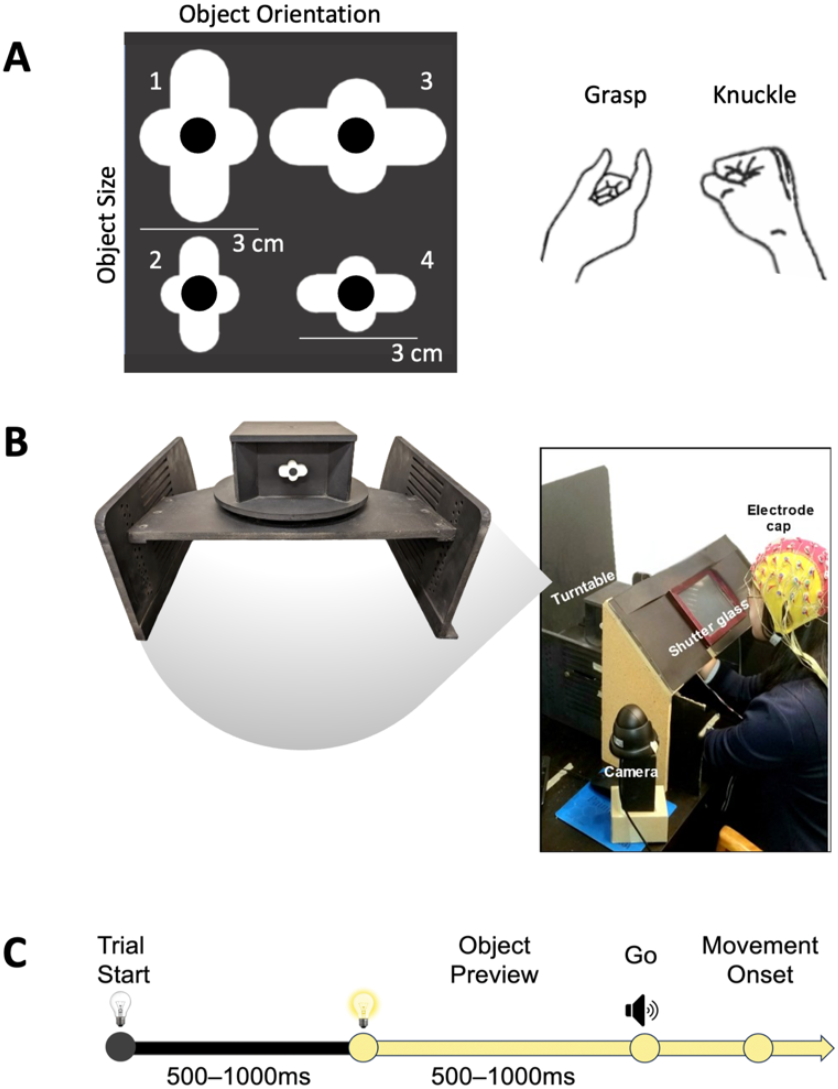
Experimental methods. **(A)** Objects and action conditions. **(B)** Experimental setup. **(C)** Timeline of a trial.

Participants always grasped along the horizontal axis of the object with their thumb and index finger of their right hand, thus the grip axis was (depending on object orientation) 67.5 mm or 45.0 mm for the larger objects, and 45.0 mm or 30.0 mm for the smaller objects. The stimuli were designed such that the grip size, but not visual size would be shared between two of the conditions, allowing for further analyses to disentangle these forms of representation.

A laser pointer was affixed to the same mount as the shutter glass and pointed onto the centre of the object. Participants were instructed to keep their eyes on the fixation point when it became visible during the trial. They were also given ear plugs to minimize the sound of the motorized turntable and thus any information about the upcoming objects that might be gleaned from the duration of the rotations.

### Experimental Design

The experimenter sat outside the room where the trial conditions were displayed to them on a monitor. There was also a camera with live feed from inside the room that allowed the experimenter to monitor the stimuli and the participant’s actions. The experimenter continuously monitored the trial from outside the room and any invalid trials (e.g. wrong actions or unstable grasps) would be manually marked.

For each trial the participant started with their hand on a button box. The turntable rotated to the appropriate object while the shutter glass was set to opaque. Furthermore, white noise played through a speaker to minimize participants’ ability to guess the upcoming object based on any auditory cues coming from the motor. This white noise always lasted three seconds, regardless of the actual duration of the rotation. To further mask the object information provided by the rotations, the rotations were varied such that, for example, to move to an object in the frame clockwise of the current object, the turntable could either rotate 90 degrees clockwise or 270 degrees counter-clockwise. Within all rotation conditions (i.e. rotation to the same object, neighbouring object, or backside object) the rotation averaged 180 degrees.

After the experimenter confirmed the object condition on the monitor matched the object displayed on the camera feed, the experimenter pressed a key to initiate the trial (Fig. 1C) beginning with the shutter glass turning transparent and revealing the laser point projected onto the black dot of the otherwise invisible object for the participant to fixate. Five hundred to 1000 ms later, LEDs turned on to illuminate the workspace for a preview of the object. After another 500-1000 ms, a beep was played as the Go signal after which participants were instructed to reach under the shutter glass and, in separate blocks of trials, either grasp the object horizontally with thumb and index finger or to touch the black dot in the middle of the object with the knuckle of their index finger while making a fist. Objects were attached to the mount via a slidable shaft, as such, participants pulled the objects roughly 2 cm out and then pushed the object back into its original place, so it was flush with the mount once again. The duration from the Go signal to when the participant lifted their hand off the button box was recorded as the reaction time. The trial ended when participants returned their hand to the button box. The LEDs then turned off and the turntable rotated to the next object.

Prior to beginning the experiment, participants completed a practice block of 10 trials to familiarize themselves with the task. Participants were told at the beginning of each experimental block what action to perform, either grasping or knuckling, and completed the same action for the duration of the block. Action blocks were randomly assigned across 24 blocks. Each block consisted of 56 trials that were randomized such that each object was displayed 14 times during each block (i.e. 2 object orientations x 2 object sizes x 14 repetitions). The experiment was completed across two approximately 3-hour long sessions, each with 12 blocks, on two separate days for each participant.

### EEG Acquisition and Preprocessing

We recorded EEG data using a 64-electrode BioSemi ActiveTwo recording system (BioSemi B.V., Amsterdam, Netherlands), digitized at a rate of 512 Hz with 24-bit A/D conversion, and the electrodes were arranged in the International 10/20 System. The electrode offset was kept below 40 mV throughout the experiment.

EEG preprocessing was performed offline in MATLAB using EEGLAB Toolbox (Delorme and Makeig, 2004) and ERPLAB Toolbox (Lopez-Calderon and Luck, 2014). In detail, signals were bandpass filtered (noncausal, 2^nd^ order Butterworth impulse response function, 12 dB/oct roll-off) with half-amplitude cut-offs at 0.1 and 40 Hz. Noisy electrodes which correlated <0.6 with nearby electrodes were interpolated (7 electrodes per subject on average), and all electrodes were then re-referenced to the average of all electrodes. Further, independent component analysis (ICA) was conducted on blocked data for each participant to identify and remove components that were associated with blinks (Jung et al., 2000) and eye movements (Chaumon et al., 2015; Drisdelle et al., 2017). The ICA-corrected data were then segmented relative to the onset of Preview (−100 to 500 ms) and Movement onset (MoveOn). There was no measurement of movement time in this study, thus -307 to 900 ms was chosen as an approximation consistent with our previous experiment (Lee et al., 2024). Lastly, invalid trials and epochs containing irregular reaction times (less than 150 ms or greater than 800 ms) were removed. As a result, an average of 1.6% of trials from each participant were discarded.

### Pattern Classification of Signals Across Time

We averaged epochs of data into ERP traces to improve the signal-to-noise ratio of spatiotemporal patterns. To do this, all blocks of the same action were pooled together (i.e. all grasping or all knuckling). Up to 14 epochs within a given block that corresponded to the same object (e.g. vertical large object) were averaged. This method resulted in 12 separate ERP traces per condition (e.g. grasping vertical large object) for Preview and Movement onset, respectively. Next, to perform multivariate noise normalization on these ERPs we calculated the covariance matrix for all time points of an epoch separately within each condition, then averaged these covariance matrices across time points and conditions (Guggenmos et al., 2018). To curb the impact of outliers on SVM-based pattern classification while keeping the number of features consistent, we z-scored traces across times and electrodes, thresholding outliers at 3 SD from the mean (for similar approaches see Nemrodov et al., 2017; Nemrodov et al., 2018).

We then divided ERP traces into temporal windows with 5 consecutive bins (5 bins * 1.95 ms ≈ 10 ms) to increase the robustness of pattern analyses. For each time bin, data from all 64 electrodes were concatenated to create 320 features which allowed for classification across time, window by window.

We performed pairwise discrimination of Object Orientation, Object Size, and Action using linear support vector machines (SVM; c = 1; LibSVM 3.22, Chang & Lin, 2011) and leave-one-out cross-validation. Cross-validation was done iteratively such that all data were at one point training and testing data. More specifically, for Action, 47 of 48 pairs of observations were used for training and one pair was used for testing and for Object Orientation and Object Size, 23 of 24 pairs of observations were used for training while one pair was used for testing. Except for Action, these analyses were performed separately on grasping and knuckling data (Action classifiers discriminated between grasping and knuckling classes).

Further, we investigated integrated representation between the visual features: Object Orientation ∩ Object Size. Again, we analysed data separately for grasping and knuckling, but this time we performed pairwise discrimination for integrated representations with logic in line with Lee and colleagues (2024; see integrated representations in “Pattern Classification of Signals Across Time”). That is, first SVMs were discouraged from learning markers of only Object Orientation or Object Size. Then, to test whether true integration occurred, the resulting SVMs were tested against an additive model of Object Orientation and Object Size (i.e., we tested whether for two features A and B the A∩B classifier outperformed the A classifier + the B classifier as a sign of true integration, e.g., Pollmann et al., 2014).

For the first step, we trained an SVM to distinguish “large vertical” objects (class 1) from “large horizontal” and “small vertical” objects (class 2). The class 1 object shares a size (“large”) with one class 2 object, and an orientation (“vertical) with the other class 2 object. Importantly, the SVM cannot perform well by learning *only* orientation or size. Because class 2 contains twice the amount of training observations compared to class 1, the cost parameter for class 1 was double that of class 2 (c = 2 vs. c = 1; Batuwita & Palade, 2013) to circumvent classifier bias towards the majority class 2. We conducted leave-one-out cross-validation such that the SVMs were trained on 33 ERP traces (11 of each of the three objects) and tested on 2 traces (the testing contained the class 1 object and one class 2 object to again, discourage skew towards class 2 which would be the case if all three objects were included in testing at each iteration). These same steps were repeated with the other three permutations of comparisons, namely: “large horizontal” vs. “large vertical” and “small horizontal”, “small vertical” vs. “large vertical” and “small horizontal”, and “small horizontal” vs. “small vertical” and “large horizontal”, and an average performance was calculated from these four analyses for the integrated representation of Orientation ∩ Object Size for each participant. For the second step, for grasping and knuckling separately we modelled an estimate of how two single-feature classifiers perform optimally together based on the representations of the Object Orientation and Object Size (i.e. the first two rows of Fig. 2; refer to integrated classification in “Pattern Classification of ERP Signals Across Time” in Lee et al., 2024 for equation and further details).

**Figure 2.**
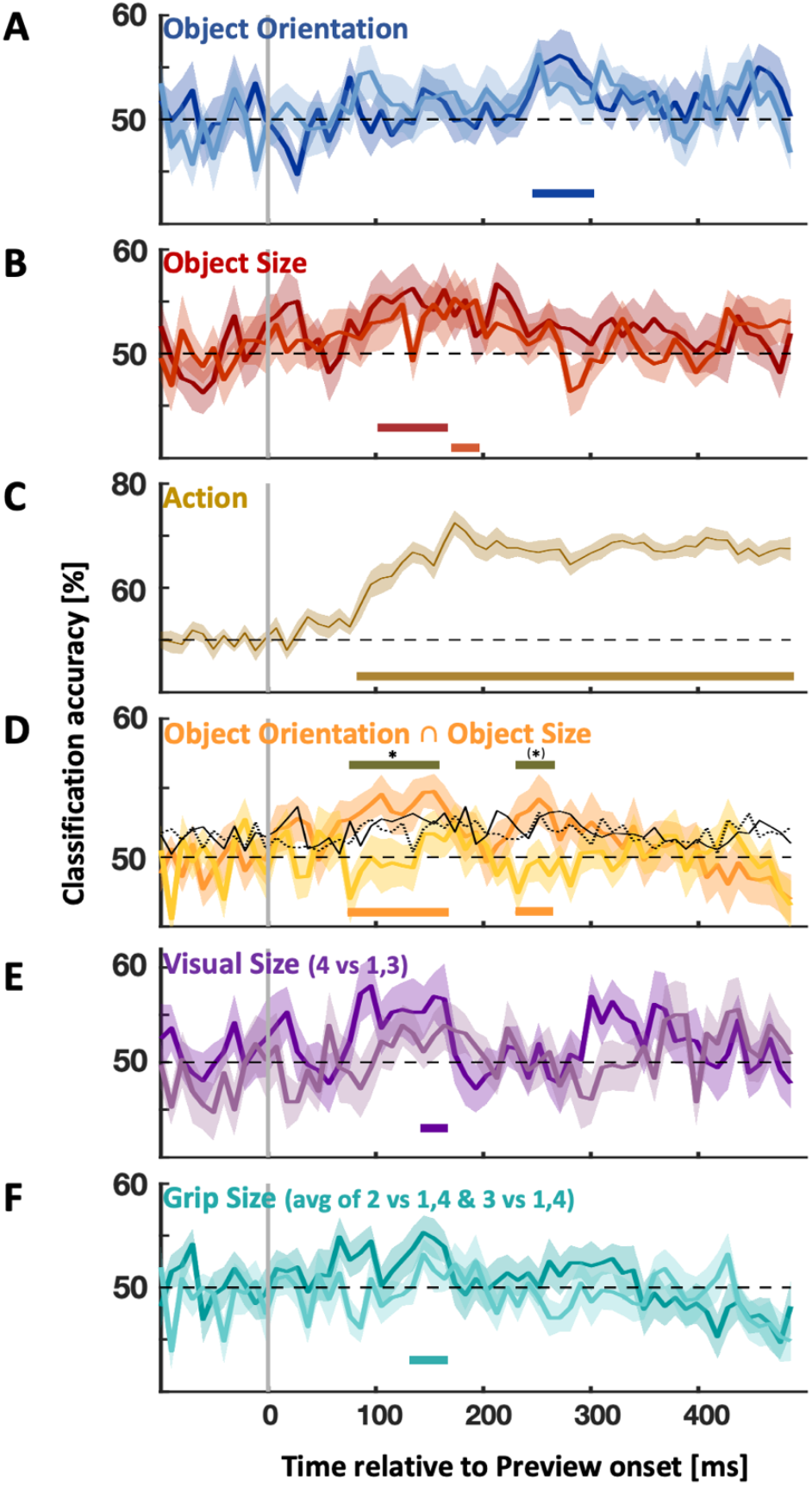
Time-resolved classification accuracy of effects aligned to Preview onset. Darker colours in each plot (i.e. curves and horizontal bars) represent grasping, lighter represent knuckling. Horizontal coloured bars denote significantly above chance classification (cluster-corrected t-test, one-tailed; p<0.05). Horizonal grey bars denote significantly better classification accuracies for grasping compared to knuckling (cluster-corrected t-test, one-tailed; p<0.05). Asterisks above bars indicate significantly higher values for integrated representations (dark orange) compared to the additive model (solid black line; two-tailed; p<0.05).

We also conducted classification for Visual Size and Grip Size, which were conditions that had an unequal number of observations for two categories of classes. Visual Size, for example, was a comparison between object 4 (Fig. 1A, the “horizontal small” object) as class 1, and objects 1 (“vertical large”) and 3 (“horizontal large”) as class 2. In this case, object 4 shares an object orientation with object 3, and a grip size with object 1. The only feature not shared between object 4 and these other two objects is the pure visual size. In this way, the analysis was set up to encourage the classifier to use visual size to discriminate between these classes (n.b., object 2 was not included in the analysis because it differed from the other objects along more than one feature dimension so that classification would have been confounded). For Grip Size, we took the average performance of two classifiers. The first was a comparison between object 2 (“vertical small” object) as class 1 versus object 1 (“vertical large” object) and 4 (“horizontal small” object) as class 2. The second classifier was between object 3 “(horizontal large” object) as class 1 versus object 1 (“vertical large” object) and 4 (“horizontal small” object) as class 2. In both instances, class 1 shares the object orientation with one object from class 2, and object size with the other object from class 2. This was to encourage classifiers to learn grip size information that is separate from object size and orientation.

Because for all classifications, class 2 contained twice as many trials, as described previously, it was necessary to set up classification analyses such that the cost parameter of the support vector was double the value the minority class 1 compared to majority class 2 (c = 2 vs. c = 1; Batuwita & Palade, 2013). We performed leave-one-out cross-validation, such that although training data had an unequal number of instances for each class, testing had one of each class (33 observations used for training and 2 for testing, where half of the testing data were from minority class 1 and half were from majority class 2) to further discourage the classifier from using only one of the two features.

For all analyses, decoding was performed for each subject separately and then averaged across participants. Further, for all analyses, data were aligned to stimulus onset (“Preview onset”).

### Statistical Analyses

The alpha level was set at .05 for all statistical tests. For behavioural data, we performed a three-way repeated measures ANOVA (Action x Object Orientation x Object Size) to test for differences between conditions in reaction time (RT). We adjusted for sphericity violations using Greenhouse-Geisser (GHG) corrections and used repeated measures t tests with Bonferroni correction for multiple comparisons for post hoc analyses.

In the case of EEG and EMG data, we assessed statistical significance using a non-parametric, cluster-based approach to determine clusters of timepoints where there were significant effects at the group level (Nichols & Holmes, 2002). We defined clusters as consecutive time points that were equivalent or exceeded the 95^th^ percentile of the distribution of t-values at each time point attained using sign-permutation tests computed 10000 times (equivalent to p<0.05, one-tailed). Next, to determine the size criteria of what constituted a significant cluster, we selected the 95^th^ percentile of maximum cluster sizes across all permutations (equivalent to p<0.05, one-tailed).

## Results

### Behavioural Results

The average reaction time (RT; time between Go onset and Movement onset) was 382 ms (SD = 138 ms). RTs submitted to a three-way repeated-measures ANOVA (Action x Object Orientation x Object Size) did not produce any significant main effects or interactions (F’s < 31.737, p’s > 0.209).

### Classification of ERPs Aligned to Preview Onset

We obtained the time-resolved classification accuracy of ERPs aligned to Preview for main effects (Object Orientation, Object Size and Action), as well as special isolated effects (Visual Size and Grip Size). In detail, Grasping Object Orientation classification was significant from 250 ms to 300 ms. Knuckling Object Orientation classification did not reach significance (Fig. 2A). Conversely, Grasping Object Size classification was significant earlier, from 105 ms to 165 ms, and there was a transient period of Knuckling Object Size significance from 175-195 ms (Fig. 2B). There were no differences between Grasping and Knuckling classification accuracy for Object Orientation or Object Size.

For Action (i.e. discriminating between Grasping and Knuckling), classification accuracy gradually rose, becoming significant starting at 85 ms and remaining robustly significant for the remainder of Preview (Fig. 2C).

Crucially though, the integrated representation of these features during Grasping significantly outperformed Knuckling from 75-155 ms and from 230-260 ms (outperformed chance from 75-165 ms and 230-260 ms; Fig. 2D, dark vs. light orange curves). To compare the integrated representation for Grasping with an additive model of separate object orientation and size representations (Fig. 2D, solid black line; see Lee et al., 2024, “Pattern Classification of ERP Signals Across Time” for equations and descriptions) with the Grasping integrated representation of object orientation and object size (Fig. 2D, dark orange curve), we took the classification accuracies of each respective curve for 75-155 ms and 230-260 ms as an a priori time frame for when to expect superadditivity. We then calculated a mean value for each of the curves during these two time periods. A paired samples t-test indicated that for the first segment of time, the Grasping integrated representations (*t*(8) = 4.39, *p* = 0.002), whereas for the second time segment the t-test just fell short of significance (*t*(3) = 3.16, *p* = 0.05).

Representations for pure Visual Size and pure Grip size for Grasping were significant from 145-165 ms (Fig. 2E) and similarly from 135-165 ms (Fig. 2F), respectively. Interestingly, these times also overlapped with significant Object Size representations and integrated Size by Orientation representations for Grasping. Neither Visual Size, nor Grip Size classification was significant for Knuckling, and there were no differences between Grasping and Knuckling classification accuracy for Visual Size or Grip Size.

## Discussion

In this study, we used time-resolved classification of EEG data recorded from participants as they grasped objects or reached and touched them with their knuckle. These objects differed in size, and they were either vertically or horizontally oriented. We investigated how the representations of these features might be modulated by action intention, in particular, whether the intention to grasp, but not the intention to reach for the objects yielded integrated representations of size and orientation. As expected, we found evidence for an integrated representation of object shape and object orientation to occur in the planning phase of a grasp.

In more detail, we found a superadditive representation of the features object size together with object orientation from 75-155 ms and trend-wise from 230-260 ms with the first segment of significance overlapping with the significant times for the Grasping representations of Visual Size (145-165 ms) and Grip Size (135-165 ms). This is consistent with the idea that the integration is related to the visual analysis of geometric properties necessary to determine stable grasp points on the surface of the object (Blake, 1995; Lederman & Wing, 2003; Kumra et al., 2022). Consistent with these putatively visual processes we found that pure representations of visual and grip size formed at about the same time. What is more, the integrated representation of size and orientation coincided with the integrated representation of size and shape that Guo and Niemeier (2024) reported to occur 140-259 ms after object onset, thereby confirming those earlier findings.

These integrated representations of object orientation and size might involve the anterior intraparietal sulcus (aIPs) which has previously been implicated in computing the geometric features of objects (Murata et al., 2000; Schaffelhofer and Scherberger, 2016). Further, deactivation of aIPS via TMS has been shown to negate the increase of orientation sensitivity typically seen when preparing to grasp over preparing to reach possibly through disrupting top-down feedback to early visual areas (Gutteling et al., 2013).

Crucially however, the present study is the first to show that integrated representations of task-relevant features size and orientation indeed arise as a function of action intention; we observed the integration only when participants intended to grasp an object, not when they merely planned to touch the object with their knuckle.

Regardless of task, participants were required to direct their attention towards the object. Thus, it is noteworthy that theories of object-based attention suggest that directing attention to an object, regardless of task, is sufficient for the integration of its features (Kahneman & Treisman, 1984; Kahneman et al., 1992; Duncan, 1984). However, our findings are clearly at odds with that prediction. Instead, the observation of task-specific integration of object features is consistent with the broader view of sensorimotor control as a nonlinear process that affords the incorporation of multiple parameters or features for its computations depending on task.

Together, this study shows that the brain integrates superadditive representations of object orientation and object size shortly after object onset. Indeed, integration likely occurs to address the computational needs of nonlinear motor control necessary for grasping. This integration is visual, reflecting the geometric analysis that is carried out, only when intending to grasp an object. Our results offer more nuance to the understanding of preparatory mechanisms underlying grasp control.

## Acknowledgement

This research was supported in part by a grant from the Natural Sciences and Engineering Research Council of Canada (NSERC).

